# European taurine vs indian indicine cattle: a comparative genomic study reveals regions of differentiation during evolution and selection during crossbreeding

**DOI:** 10.64898/2026.02.05.704058

**Authors:** Poorvishaa V Muthusamy, Prerna Gupta, Nupur Upadhyay, Rajesh Vakayil Mani, Manpreet Kaur, Balu Bhaskar, Rajeev Raghavan Pillai, Thankappan Sajeev Kumar, Thapasimuthu Vijayamma Anilkumar, Sidharth Kulkurni, Sarwar Azam, N. Sadananda Singh

## Abstract

Cattle are broadly classified into two subspecies: *Bos taurus*, adapted to temperate climates, and *Bos indicus*, adapted to tropical environments. Indicine cattle show better heat tolerance and stronger disease resistance, whereas taurine cattle are known for higher milk yield and better meat quality. Improving milk yield while maintaining resistance to heat stress and infectious diseases is an important objective in cattle breeding. Marker-assisted selection is an effective approach for improving economically important traits, highlighting the need to understand genetic differences between taurine and indicine cattle. However, genome-wide comparisons between European taurine and Indian indicine cattle remain limited. To address this gap, whole-genome sequencing of 48 Indian indicine cattle was performed and combined with publicly available data. This enabled a comparative genomic analysis of 74 Indian indicine and 83 European taurine individuals to identify genes involved in heat tolerance and immune response. Genome-wide analyses using Fst and XP-CLR identified 4,343 and 1,457 differentiated genes, respectively, with 826 genes common to both methods. These genes were mainly associated with immune response, protein stability, and cytoskeletal structure. Strong selection signals were observed in three heat shock protein genes (*DNAJC11*, *DNAJC5*, and *DNAJB11*) and 229 immune-related genes. To examine the inheritance of these genes through crossbreeding, a haplotype-resolved genome assembly was generated for the Indian crossbreed Sunandini, which showed predominantly taurine ancestry (69.13-96.04%), with a smaller indicine contribution (1.22-1.87%). Several genes related to heat tolerance and immune response were inherited exclusively from indicine cattle, highlighting their importance for environmental adaptation and future breeding programs.

## Introduction

Cattle play an important role in human society by supporting national economies, rural livelihoods, and food and nutritional security. Long-term domestication and management in different environments led to the divergence of two major cattle subspecies: *Bos taurus taurus* (commonly referred to as *Bos taurus*), adapted to temperate climates, and *Bos taurus indicus* (commonly referred to as *Bos indicus*), adapted to tropical environments (Pitt et al. 2018; Arbuckle and Kassebaum 2021). Environmental factors such as temperature, feed availability, and disease pressure acted as strong selective forces, resulting in clear morphological, physiological, and cellular differences between the two subspecies (Hansen 2004; Cooke et al. 2020; Okamoto et al. 2025).

*Bos indicus* cattle evolved under prolonged exposure to high temperature, humidity, limited nutrition, and heavy pathogen load. As a result, they show strong heat tolerance, characterised by smoother, less dense hair coats, loose skin, a higher density of sweat glands located close to the skin surface, smaller metabolically active organs, and lower basal metabolic rates. These traits improve heat dissipation and reduce internal heat production, allowing indicine cattle to maintain body condition under heat stress and poor nutrition. In contrast, *Bos taurus* cattle have denser hair coats, tighter skin, fewer sweat glands, and higher metabolic heat production, which makes them more susceptible to heat stress (Hansen 2004; Jian et al. 2014; Scheffler 2022; Okamoto et al. 2025). Experimental studies have further shown that elevated temperatures more severely affect embryonic development and immune cell survival in taurine cattle, with reduced blastocyst development and lymphocyte viability compared with indicine cattle. Together, these findings indicate greater cellular heat tolerance in *Bos indicus* (Paula-Lopes et al. 2003; Hansen 2004).

In addition to heat tolerance, *Bos indicus* cattle also show greater resistance to infectious and parasitic diseases, even when maintained under the same environmental conditions as taurine cattle. They exhibit partial resistance to several tick-borne diseases, including babesiosis (*Babesia bovis* and *B. bigemina*), anaplasmosis (*Anaplasma marginale*), tropical theileriosis (*Theileria annulata*), and East Coast fever (*Theileria parva*) (Piper et al. 2009; Glass et al. 2005; Gul et al. 2015; Tabor et al. 2017; Jonsson et al. 2008). Studies on *Rhipicephalus (Boophilus) microplus* infection further demonstrate that indicine breeds such as Brahman mount more regulated T-cell–mediated adaptive immune responses and maintain balanced cytokine production. In contrast, taurine breeds such as Holstein-Friesians show stronger innate inflammatory responses, marked by elevated pro-inflammatory cytokines. By limiting excessive inflammation while maintaining effective adaptive immunity, indicine cattle experience milder disease outcomes, highlighting genetic differences in immune regulation between the two subspecies (Piper et al. 2009; Glass et al. 2005).

Despite these adaptive advantages, *Bos indicus* cattle generally produce much less milk than *Bos taurus* cattle. In indicine-dominated regions such as India, Pakistan, and much of Africa, average milk yield is approximately 500 kg per cow per year, whereas taurine-dominated regions in North America and Europe achieve yields exceeding 5,000 kg per cow per year. This large productivity gap contributes to low per capita milk availability in many tropical regions (Cunningham and Syrstad 1987). To address this limitation, high-yielding European *Bos taurus* breeds were introduced into tropical production systems. However, these efforts were largely unsuccessful because taurine cattle are poorly adapted to hot and humid environments. Chronic heat stress reduces feed intake and metabolic activity, leading to negative energy balance and milk yield losses of 25–40%, while also increasing disease susceptibility. As a result, pure taurine breeds are generally unsuitable for sustainable dairy production in the tropics (Sesay 2023; Kumar et al. 2020; Habimana et al. 2023; Abdel Rahman and Alemam 2008).

Crossbreeding between indicine and taurine cattle has therefore been widely adopted to combine higher milk yield with environmental adaptation. Although initial productivity gains have been achieved, long-term outcomes are often inconsistent. The development of stable crossbred populations typically requires two to three decades of controlled breeding and high management inputs, and only a few crossbred populations have successfully adapted to tropical conditions (Wakchaure et al. 2015; Mandal et al. 2013; Pandey 2020). Among these, Sunandini cattle represent one of the most successful crossbred dairy populations in India. Sunandini was developed under the Indo–Swiss Project in Kerala in 1963 by crossing local non-descript cattle with taurine breeds such as Brown Swiss, Holstein Friesian, and Jersey, followed by nearly three decades of selective breeding and optimization (Chacko 1994; Taneja 2000). According to the 2012 Livestock Census, Kerala had 13.29 lakh cattle, of which 12.52 lakh (approximately 94%) were Sunandini. Consequently, Sunandini cattle contribute about 85–90% of the state’s total milk production, with crossbred cows producing an average of 10.79 kg of milk per day. (Government of Kerala 2018; Kerala Livestock Development Board 2012) Despite their high productivity and economic importance, Sunandini cattle remain vulnerable to tropical diseases and reproductive disorders, and require higher management inputs than indigenous breeds. These limitations suggest that conventional crossbreeding alone may not be sufficient to ensure long-term sustainability (Shivakumara et al. 2018; Elayadeth-Meethal and Kolathingal-Thodika 2025).

Recent advances in genomics provide new opportunities to improve cattle breeding by enabling the identification of genes associated with milk yield, heat tolerance, and disease resistance. However, the effective application of genomic selection requires a clear understanding of the genetic differences between *Bos indicus* and *Bos taurus*, as well as insight into how adaptive genomic regions are inherited and maintained in crossbred populations. To address this limitation, a comparative genomic analysis of *Bos indicus* and *Bos taurus* cattle was conducted to identify the genomic regions associated with heat shock tolerance and immune response. Whole-genome sequencing of 48 Indian indicine cattle was performed to detect the highly differentiated regions between indicine and taurine populations. These regions were further examined in a tropically adapted crossbred population using Sunandini cattle as a model. In addition, a high-quality Sunandini reference genome was generated using a combination of long- and short-read sequencing technologies to assess the relative contributions of indicine and aurine ancestry to traits related to milk production, heat tolerance, and immunity.

## Results

### Whole-genome sequencing of *Bos indicus* breeds

High-quality whole-genome sequencing data were generated for three *Bos indicus* breeds: 23 Sahiwal, 22 Gir, and 3 Vechur to investigate genetic differentiation between *Bos taurus* and *Bos indicus.* The sequencing libraries consisted of 150 bp paired-end reads, with individual samples yielding between 665 million and 1,448 million reads for Sahiwal, 747 million to 1,510 million reads for Gir, and 865 million to 1,118 million reads for Vechur before quality filtering. These translated into base counts ranging from 99 Gbp to 217 Gbp (Sahiwal), 112 Gbp to 226 Gbp (Gir), and 130 Gbp to 169 Gbp (Vechur). The GC content prior to filtering ranged from 43.35% to 49.11% across samples. After quality filtering, the datasets retained between 578 million and 1,346 million reads for Sahiwal, 642 million to 1,015 million reads for Gir, and 728 million to 856 million reads for Vechur, corresponding to effective base counts of 77-185 Gbp (Sahiwal), 77-138 Gbp (Gir), and 117-129 Gbp (Vechur). The GC content remained stable (41.5%-48.8%) after filtering (Table 1). Mapping efficiency to the ARS-UCD1.2 reference genome was high, with most samples achieving over 99% alignment, except for a single Gir sample that showed a lower mapping rate (66.37%). The number of SNPs/variants range from 15,690,953 to 17,898,308 in the newly resequenced genomes. The newly resequenced *Bos indicus* datasets were subsequently combined with publicly available *Bos taurus* and *Bos indicus* genomes obtained from the Bovine Genome Variation Database and the Selective Signatures Database, representing diverse European *Bos taurus* and Indian *Bos indicus* populations. The final dataset consists of 74 *Bos indicus* and 83 *Bos taurus* individuals for downstream comparative genomic analyses.

**Table 1.** Summary of whole genome sequencing read statistics.

| Breed | No. of Samples | Read Length (bp) | Read Count (M) Before Filtering (Mean ± SD) | Base Count (G) Before Filtering (Mean ± SD) | GC Content (%) Before Filtering (Mean ± SD) | Read Count (M) After Filtering (Mean ± SD) | Base Count (G) After Filtering (Mean ± SD) | GC Content (%) After Filtering (Mean ± SD) | Mapping (%) (Mean ± SD) |
| --- | --- | --- | --- | --- | --- | --- | --- | --- | --- |
| Sahiwal | 22 | 150 | 872.62 ± 168.71 | 130.89 ± 25.30 | 45.63 ± 0.88 | 773.76 ± 161.77 | 108.78 ± 22.35 | 45.33 ± 0.89 | 99.47 ± 0.26 |
| Gir | 23 | 150 | 890.77 ± 161.06 | 133.61 ± 24.16 | 47.02 ± 1.34 | 780.30 ± 113.11 | 107.30 ± 15.65 | 46.58 ± 1.50 | 98.16 ± 6.90 |
| Vechur | 3 | 150 | 961.10 ± 137.54 | 145.12 ± 20.76 | 44.18 ± 0.13 | 781.30 ± 113.11 | 123.30 ± 5.95 | 44 ± 0.53 | 99.56 ± 0.33 |

### Principal component analysis and chromosome-level synteny analysis of *Bos indicus* and *Bos taurus*

Principal component analysis o*f Bos indicus, Bos taurus*, and other cattle genome-wide SNP data revealed three main clusters: Indian *Bos indicus*, Chinese *Bos indicus*, and European *Bos taurus* (Supplemental Figure S1). Based on this separation, subsequent analyses focused on comparisons between Indian *Bos indicus* and European *Bos taurus*. To examine whether genetic differences between these two groups are associated with large-scale chromosomal changes, a macrosynteny analysis was performed to assess conservation of gene order. Chromosome-level genome assemblies were obtained from NCBI, including four *Bos taurus* breeds (Charolais, Hereford, Holstein-Friesian, and Jersey) and three *Bos indicus* genomes, representing three Indian breeds (Nellore, Vechur, and Tharparkar).

Initial synteny maps generated using the default Chordate-Wide ODP database were highly noisy and lacked clear linkage groups, limiting their interpretation. To improve resolution, a cattle-specific synteny database was constructed using Banteng as a wild outgroup, Hereford as the *Bos taurus* representative, and Nellore as the *Bos indicus* representative. Synteny analysis using this custom database showed strong macrosyntenic conservation between *Bos taurus* and *Bos indicus* genomes, with no evidence of large-scale chromosomal rearrangements (Supplemental Figure S2). These results indicate that differentiation of *Bos indicus* and *Bos taurus* being a result of sequence-level variation rather than chromosomal structural changes.

### Fst-based single-SNP differentiation analysis of heat shock genes between *Bos taurus* and *Bos indicus*

To investigate the genetic basis of heat tolerance and immune competence differences between European *Bos taurus* and Indian *Bos indicus*, genome-wide single-nucleotide polymorphism (SNP) analyses were performed using Fst, a measure of population differentiation that highlights genomic regions under divergent selection. From a total of approximately 70.22 million SNPs, multiallelic variants were excluded using GATK SelectVariants, retaining only biallelic SNPs for further analysis. Stringent quality control measures were applied to ensure data reliability, removing SNPs with more than 20% missing genotypes and those with a minor allele frequency (MAF) below 0.05. After filtering, about 17.21 million high-quality SNPs were retained for downstream analyses, of which 1.02 million SNPs (6%) exhibited high genetic differentiation (Fst ≥ 0.8). These highly differentiated SNPs were distributed across the genome and mapped to 14,270 genes (Supplemental Figure S3).

To focus on traits directly relevant to heat adaptation, the heat shock protein (HSP) gene family was examined, which plays a crucial role in maintaining cellular protein homeostasis under thermal stress through its molecular chaperone activity. Among the 97 known HSP genes (https://www.genenames.org), 52 genes carried highly differentiated SNPs (1549 SNPs) between Indian *Bos indicus* and European *Bos taurus* (Supplemental Table S1). Manual inspection of allele frequencies confirmed consistent population-level differences between taurine and indicine cattle, supporting their potential role in thermal adaptation. Since longer genes naturally accumulate more variants and potentially bias differentiation estimates, SNP counts were normalized by gene length (variants per kilobase) (Supplemental Table S1). After normalization, several genes, including *DNAJC11*, *DNAJC8* and *HSPA4*, remained prominently differentiated, highlighting their likely contribution to the enhanced heat tolerance observed in *Bos indicus*.

Considering the environmental similarities between African and Indian cattle, both of which are routinely exposed to high temperatures and nutritional stress, the analysis was extended to include 28 African *Bos indicus* samples that formed a distinct cluster from European cattle in the PCA plot (Supplemental Figure S1). These samples were compared with 83 European *Bos taurus* individuals. Applying the same quality control filters as in the comparison with indian indicine, 1,048 highly differentiated SNPs (Fst ≥ 0.8) were identified, mapping to 37 heat shock protein (HSP) genes (Supplemental Table S2), indicating strong allele frequency differences between populations. To identify consistent signals of heat adaptation, results from the Indian vs. European and African vs. European comparisons were intersected. This overlap revealed 905 SNPs across 35 common HSP genes (Supplemental Table S3) that consistently exhibited population differentiation in both datasets, supporting their potential role in thermal and climate adaptation.

### Window-based Fst analysis reveals selection signatures in heat shock protein families between Indicine and Taurine Cattle

To refine the understanding of genomic regions associated with heat adaptation, a window-based Fst analysis was conducted between Indian *Bos indicus* and European *Bos taurus*. Unlike single-SNP analyses, this approach scans the genome using sliding windows, allowing detection of broader regions enriched with highly differentiated variants that may represent selection hotspots. Using 50 kb windows with 25 kb steps, a total of 99,339 genomic windows were analyzed, resulting in the identification of 4,750 outlier windows with Fst values greater than 0.7. The distribution of these windows across the genome is shown in the Manhattan plot (Figure 1A). These outlier windows collectively mapped to 4,343 genes, many of which were enriched in biological processes related to cellular response to stimulus (981 genes), response to stress (509 genes), programmed cell death (248 genes), and regulation of immune system processes (215 genes). The Gene Ontology (GO) cluster maps illustrating enrichment for biological processes are presented in Supplemental Figures S6.

**Figure 1.**
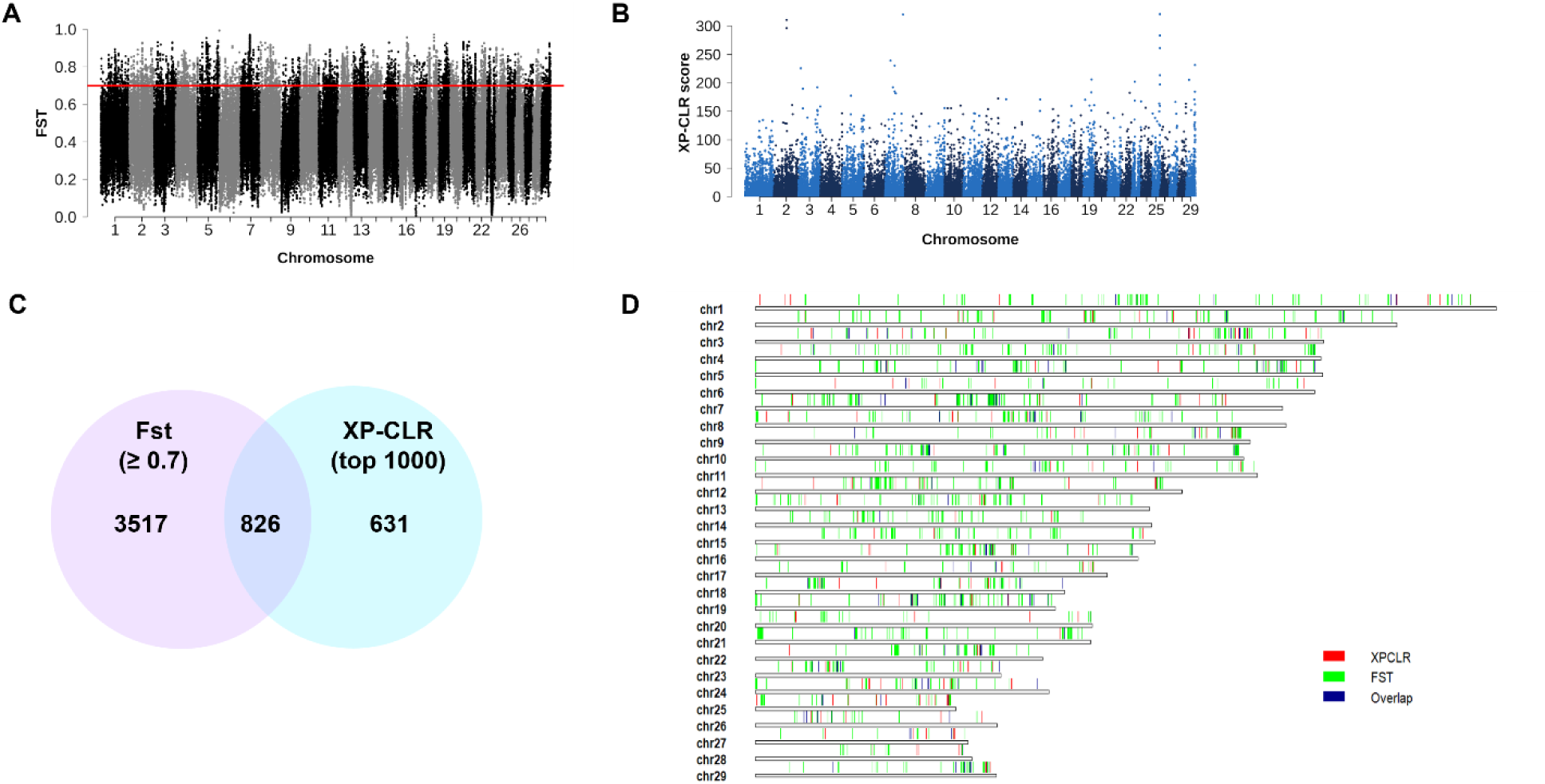
Genome-wide differentiation and selective sweep analyses between Indian *Bos indicus* and European Bos taurus cattle (A) Genome-wide window-based Fst comparison between Indian *Bos indicus* and European Bos taurus (B) Genome-wide window-based selective sweep analysis using XP-CLR between Indian *Bos indicus* and European Bos taurus (C) Venn diagram showing the overlap between genes located within regions showing Fst values ≥ 0.7 and the top 1000 XP-CLR scores (D) Karyogram illustrating genomic regions with the top 1000 XP-CLR scores (Orange), window-based Fst ≥ 0.7 (Blue), and overlapping regions identified in both analyses (Black)

When the analysis was restricted to heat shock protein (HSP) genes within these highly differentiated regions, 15 genes showed consistent and strong differentiation signals. These included 10 genes from the HSP40 (DNAJ) family (*DNAJB4, DNAJB6, DNAJB11, DNAJB12, DNAJC5, DNAJC8, DNAJC11, DNAJC17, DNAJC18* and *DNAJC19*) three genes from the HSP70 (HSPA) family (*HSPA4, HSPA9* and *HSPA12B*), and two genes from the HSP90 family (*HSP90AA1* and *HSP90AB1*). A similar window-based Fst analysis between African *Bos indicus* and European *Bos taurus* also revealed strong differentiation in a subset of HSP genes, comprising five from the HSP40 (DNAJ) family (*DNAJC5, DNAJC8, DNAJC11, DNAJC14* and *DNAJC18*) and three from the HSP70 (HSPA) family (*HSPA4, HSPA9* and *HSPA12B*). The comparative summary of HSP gene differentiation across both analyses is presented in Supplemental Table S4.

### Cross-Population Composite Likelihood Ratio (XP-CLR) analysis confirms selection signatures in heat shock protein genes of Indicine cattle

To confirm that the highly differentiated regions represent true signals of selection rather than random variation, a cross-population composite likelihood ratio (XP-CLR) analysis was performed between Indian *Bos indicus* and European *Bos taurus*. This method combines differences in allele frequencies with patterns of linkage disequilibrium to identify selective sweeps with higher resolution. The analysis was conducted using a 50-kb window size and a 25-kb step size. Among the 99,339 windows examined, 34,000 windows exhibited XP-CLR scores. The genome-wide distribution of XP-CLR signals is shown in Figure 1B. The top 1,000 windows with the highest scores were selected, corresponding to 1,457 genes. Functional enrichment analysis indicated that these genes were predominantly involved in pathways such as nucleobase-containing compound metabolic process (283 genes), regulation of response to stimulus (203 genes), protein metabolic process (178 genes), response to external biotic stimulus (69 genes), and regulation of immune system process (64 genes). The gene ontology clusters for biological process are shown in Supplemental Figures S7. Within these regions, one chaperone-related gene, *BBS12*, and four HSP40 (DNAJ) family genes, *DNAJB11, DNAJC5, DNAJC11*, and *DNAJC13*, were also detected.

A comparative analysis of the top 1,000 XP-CLR regions and genomic windows with Fst values greater than 0.7 identified 826 genes were common to both analyses, illustrated by the Venn diagram and karyograms in Figures 1C and 1D. Functional enrichment of these overlapping genes revealed significant representation in pathways related to metabolic processes, including fatty acid and amino acid metabolic processes, cell morphogenesis processes, protein stability process, endocytosis and exocytosis, regulation of immune system processes, apoptotic process and development process related to reproduction, as shown in Figure 2A. Among these, three heat shock protein genes *DNAJB11*, *DNAJC5*, and *DNAJC11* were consistently identified across both Fst and XP-CLR analyses. A comparative summary of HSP gene differentiation, including Fst and XP-CLR values, is presented in Tables 2, with detailed variant consequences for these genes provided in Table 3. Multiple variants in upstream, downstream, and UTR regions suggest that regulatory mutations may contribute to altered gene expression. These HSP40 (DNAJ) family genes are involved in protein folding, mitochondrial organization, and cellular stress response, indicating a key role in heat tolerance and environmental adaptation in *Bos indicus*. Collectively, results from single-SNP Fst, window-based Fst, and XP-CLR analyses highlight these genes as strong candidates under positive selection.

**Figure 2.**
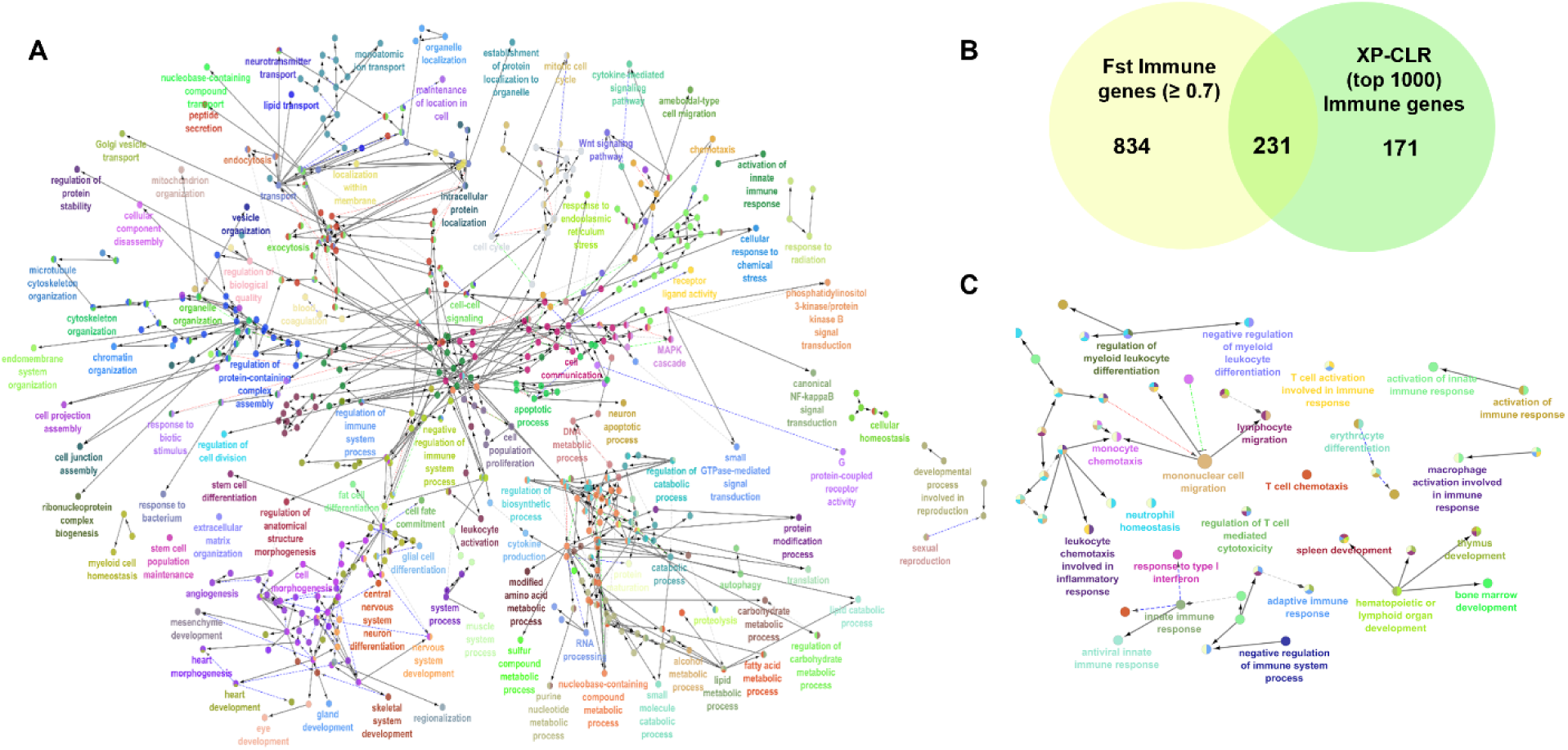
Functional enrichment of genomic regions under positive selection between Indian *Bos indicus* and European Bos taurus cattle. (A) Gene Ontology (Biological Process) enrichment of common genes identified from regions with the top 1000 XP-CLR scores and window-based Fst ≥ 0.7. (B) Venn diagram showing the overlap between immune-related genes identified from regions with Fst ≥ 0.7 and the top 1000 XP-CLR scores. (C) Gene Ontology (Immunological Process) enrichment of common genes identified from regions with the top 1000 XP-CLR scores and window-based Fst ≥ 0.7.

**Table 2.** HSP genes under positive selection identified by Fst and XP-CLR analyses with corresponding scores.

| Heat shock genes | Fst | XP-CLR |
| --- | --- | --- |
| DNAJC18 | 0.97 | 14.58 |
| DNAJC11 | 0.91 | 88.93 |
| HSPA12B | 0.86 | - |
| DNAJC5 | 0.83 | 90.07 |
| HSPA9 | 0.8 | 6.70 |
| DNAJB12 | 0.76 | 24.85 |
| HSPA4 | 0.76 | 52.01 |
| DNAJB4 | 0.75 | 11.12 |
| DNAJC17 | 0.75 | 11.54 |
| DNAJC8 | 0.75 | - |
| DNAJC19 | 0.73 | 7.59 |
| HSP90AA1 | 0.72 | 25.94 |
| HSP90AB1 | 0.71 | 27.1 |
| DNJB11 | 0.7 | 69.62 |
| DNAJB6 | 0.7 | 40.72 |
| BBS12 | 0.67 | 93.22 |
| DNAJC13 | 0.64 | 84.85 |

**Table 3.**
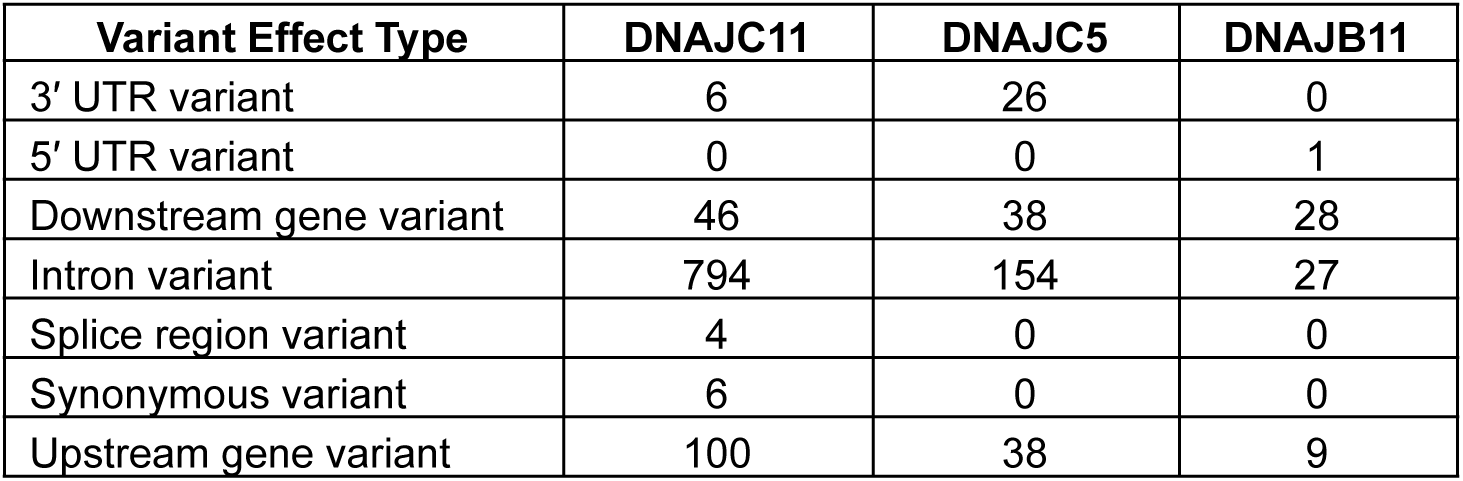
Variant consequences for common HSP genes identified in both Fst and XP-CLR analyses.

### Window-based Fst and XP-CLR analyses identify selection signatures in immune-related genes

To investigate the genetic basis of enhanced disease resistance in *Bos indicus* cattle, a comprehensive set of 7,606 immune-related genes was compiled from four major immunogenomic databases: InnateDB, ImmPort, the Immunogenetic Related Information Source (IRIS), and the Immunome Database. Intersection of these datasets with genes identified from the window-based Fst analysis revealed 1,065 immune-related genes with Fst ≥ 0.7, including 100 innate immune genes located within regions of high genetic differentiation between *Bos indicus* and *Bos taurus*. The Gene Ontology [GO] enrichment maps for immunological processes are shown in Supplemental Figures S8. Additionally, 402 immune genes, including 39 innate immune genes, were identified among the top 1,000 XP-CLR candidate regions, indicating strong differentiation. The Gene Ontology [GO] enrichment maps for immunological processes are shown in Supplemental Figures S9. A comparative analysis of Fst and XP-CLR results detected 231 immune-related genes under potential positive selection, including 24 innate immune genes (Venn diagram, Figure 2B). GO enrichment analysis of these candidate loci revealed significant representation in pathways associated with lymphocyte activation, regulation of T cell activation, leukocyte migration, monocyte chemotaxis, and response to type I interferon (Figure 2C).

Variant annotation of these immune-related genes (Table 4) revealed extensive regulatory and coding variation, including 3′ UTR (1,237), 5′ UTR (401), missense (757), synonymous (1,974), splice region (452), and stop/start-altering variants, indicating that both regulatory and coding mutations contribute to immune gene differentiation between *Bos indicus* and *Bos taurus*. The overlap between Fst and XP-CLR identified loci, combined with the variant summary, provides strong evidence that both innate and adaptive immune pathways have been targets of natural selection, highlighting enhanced immunological resilience and disease resistance in *Bos indicus* and supporting adaptation to tropical environments.

**Table 4:**
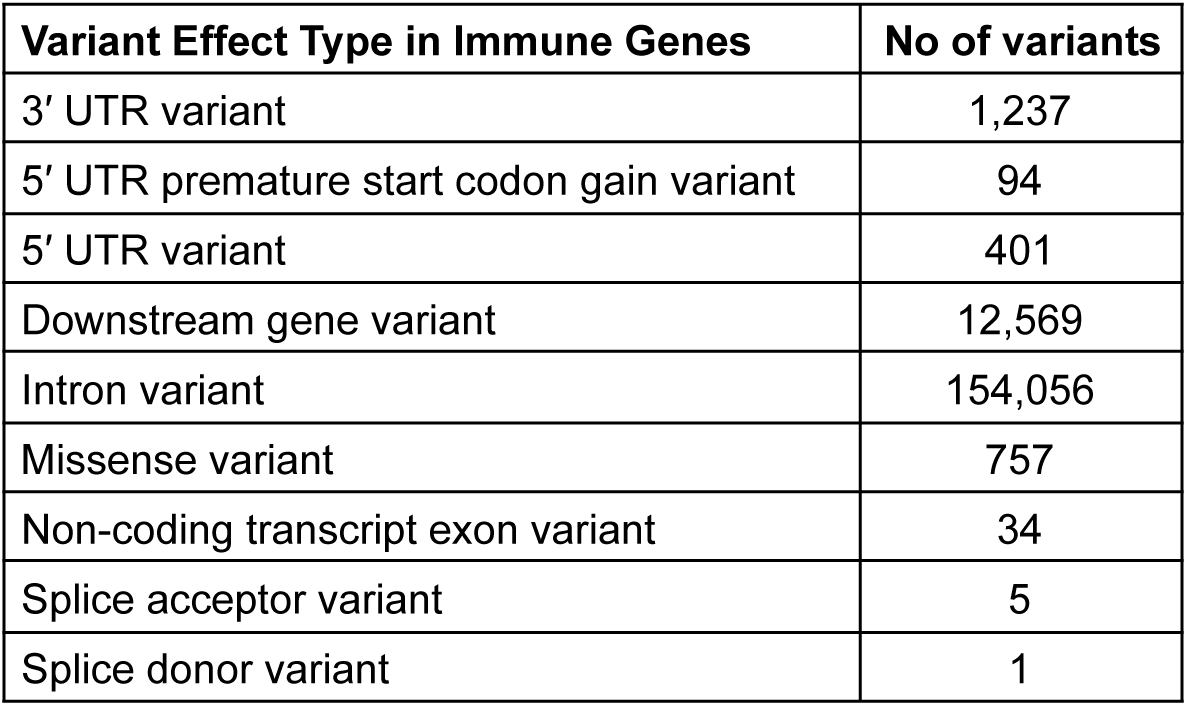
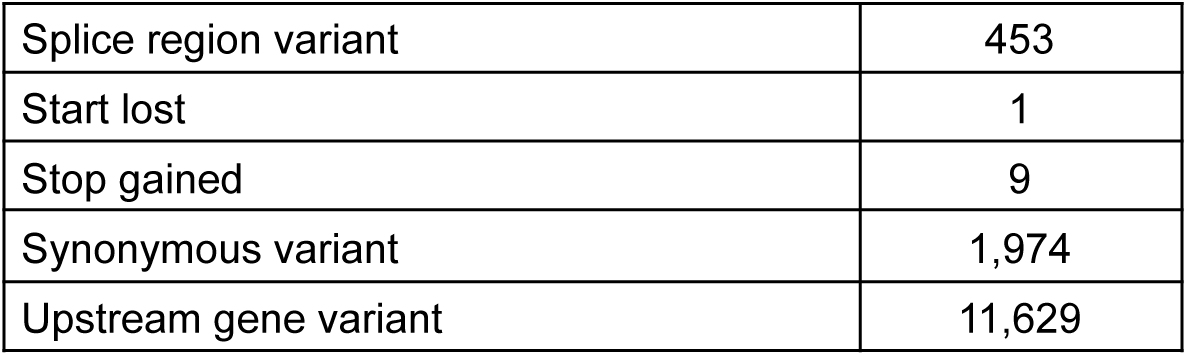
Summary of functional variant consequences detected in immune genes under positive selection.

### Hybrid genome assembly and quality assessment of the Sunandini crossbreed

To investigate whether genomic regions highly differentiated for heat tolerance and disease resistance between *Bos indicus* and *Bos taurus* have been inherited in a successfully adapted crossbreed, Sunandini cattle of Kerala were selected for analysis. This breed exhibits remarkable adaptation to tropical environments, enhanced disease resistance, and high milk productivity, making it an ideal model to study the genomic basis of successful adaptation. To uncover the underlying genomic architecture, a high-quality hybrid genome assembly was generated to identify and characterize differentiated regions contributing to adaptation and performance. Blood DNA samples were collected from five Sunandini individuals, including a family trio consisting of a dam (MT432), a sire (MT433), and their calf (MT431), along with two paternal siblings of MT431. All samples were sequenced using the Illumina short-read platform, while the calf (MT431) was additionally sequenced with Oxford Nanopore long reads to facilitate hybrid assembly. Sequencing statistics for all samples are summarized in Table 5.

**Table 5:** Summary of short-read Illumina and long-read Oxford Nanopore sequencing data.

| Short-read Illumina platform sequencing details |  |  |  |  |  |
| --- | --- | --- | --- | --- | --- |
| Sample ID | Read Length (bp) | Total reads (Forward + Reverse) | Total bases (bp) | GC (%) | Coverage (x) |
| MT429 | 151 | 90,840,8926 | 137,169,747,826 | 44.16 | 50.8x |
| MT430 |  | 98,857,1228 | 149,274,255,428 | 44.16 | 55.3x |
| MT431 |  | 89,170,0584 | 134,646,788,184 | 44.12 | 49.9x |
| MT432 |  | 86,855,3002 | 131,151,503,302 | 43.83 | 48.6x |
| MT433 |  | 99,061,7446 | 149,583,234,346 | 44.10 | 55.4x |
| Long-read Nanopore platform sequencing details |  |  |  |  |  |
| Sample ID | Read Length (bp) | Total Reads (Single end) | Total bases (bp) | GC (%) | Coverage (x) |
| MT431 | 9503 | 83,86,279 | 79,69,95,12,550 | 43.37 | 29.51 |

A hybrid de novo genome assembly was generated for the Sunandini calf (MT431) using CLC Genomics Workbench v22.0.5 with both Oxford Nanopore long reads and Illumina short reads. Illumina sequencing produced 156 GB of raw data, corresponding to 49.86× coverage, while Oxford Nanopore sequencing generated 152 GB of data (50.5× coverage). The assembly workflow was performed in sequential steps. Initially, contig-level de novo assembly was carried out using long, error-prone Nanopore reads. To improve base accuracy, the draft assembly was polished with high-quality Illumina short reads, resulting in 1,831 contigs. These contigs were subsequently scaffolded to the chromosome-level genome assembly using RagTag with the *Bos taurus* ARS-UCD1.2 genome as a reference, followed by gap patching to improve contiguity. The intermediate assembly had a total length of 2,673,071,955 bp. To further refine sequence accuracy, three additional rounds of Illumina-based polishing were performed. The final hybrid genome assembly consisted of 169 scaffolds, with a total length of 2,672,694,680 base pairs (bp) and an N50 of 102 megabases (Mb). Of these, 30 scaffolds matched the chromosome sizes of *Bos taurus*. Detailed assembly statistics are presented in Table 6, with the genome assembly visualized as a snail plot in Figure 3A

**Table 6.** Sunandini genome assembly statistics.

| Metric | Value |
| --- | --- |
| Number of scaffolds | 169 |
| Total length (bp) | 2,672,694,680 |
| Minimum scaffold length (bp) | 33,798 |
| Average scaffold length (bp). | 15,814,761.40 |
| Maximum scaffold length (bp) | 157,298,684 |
| Number of contig | 1,831 |
| Scaffold N50 (bp) | 102,000,000 |
| Percent gaps (Ns) | 0.01% |
| Complete BUSCOs (%) | 95.40% |
| Single-copy BUSCOs (%) | 93.60% |
| Duplicated BUSCOs (%) | 1.80% |
| Fragmented BUSCOs (%) | 1.50% |
| Missing BUSCOs (%) | 3.10% |
| Total BUSCO groups searched | 9,226 |

**Figure 3.**
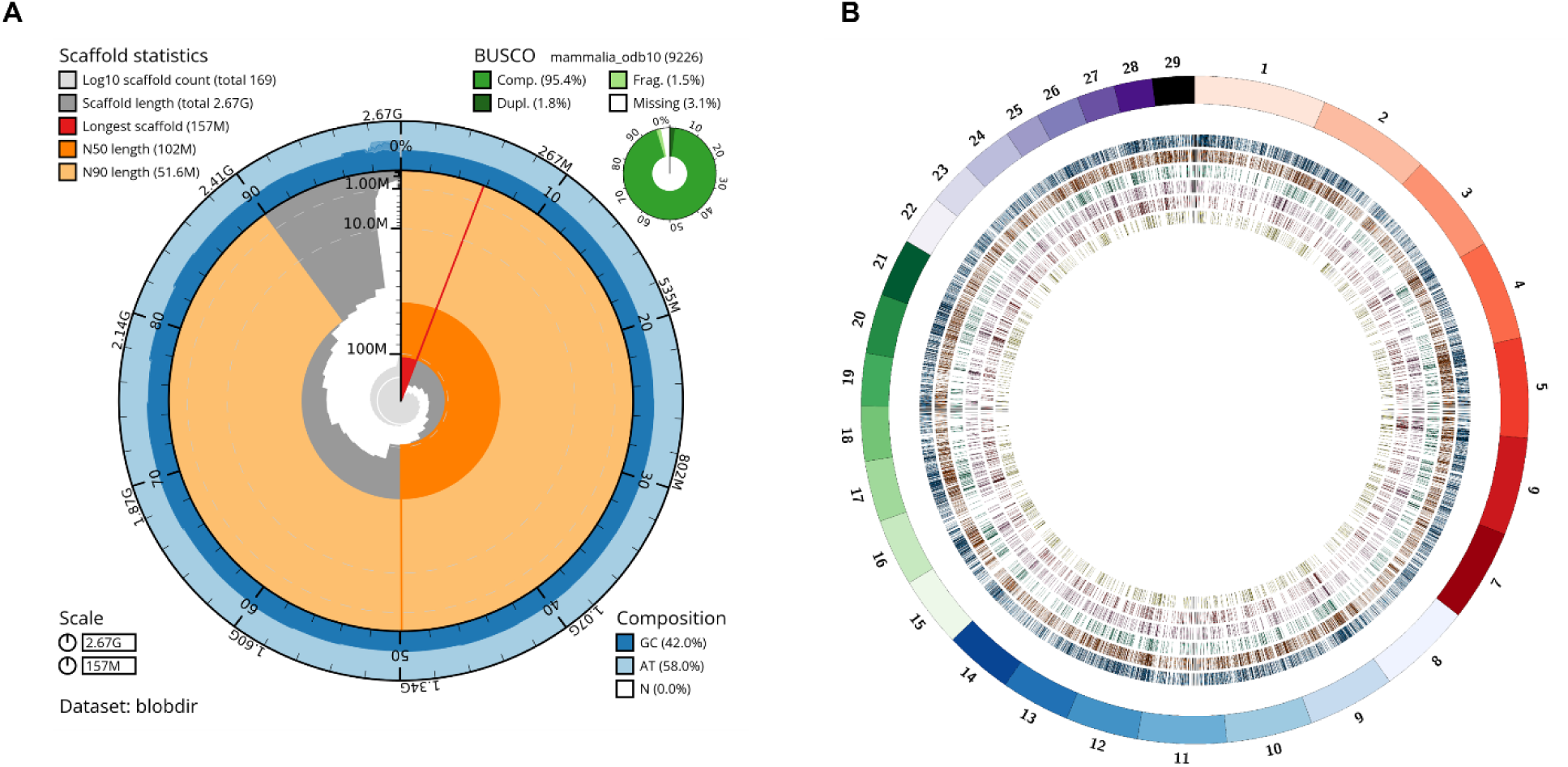
Genome assembly metrics and structural variation in Sunandini (A) Snail plot summarizing the assembly metrics of the Sunandini genome, including the length of the longest contig (157 Mb; red line), N50 (102 Mb; dark orange), N90 (51.6 Mb; light orange), and base composition, along with the BUSCO completeness assessment based on the mammalian dataset (n = 9,226).(B) Circos plot of structural variants in the Sunandini genome relative to Bos taurus ARS-UCD1.2. The outer circle shows chromosomes; inner rings represent insertions (blue), deletions (orange), repeat contractions (green), tandem expansions (pink), tandem contractions (red), and repeat expansions (yellow).

To assess potential contamination, all contigs were queried against the NCBI nucleotide (nt) database using BLASTN, and all sequences showed best hits exclusively to *Bos taurus*, confirming the absence of detectable non-bovine sequences in the assembly. Assembly quality was further evaluated by mapping all Illumina short reads back to the final genome assembly, which resulted in a 99.7% mapping rate, indicating that nearly all sequencing data were represented in the assembly. To evaluate completeness, a Benchmarking Universal Single-Copy Orthologs (BUSCO) analysis (Simao et al. 2015) was performed in genome mode using the Mammalia lineage dataset. The assembly contained 95.4% complete BUSCOs, indicating high accuracy and near-completeness of the assembled genome. A summary of BUSCO completeness metrics is provided in Table 6.

### Trio-based haplotype-resolved assembly

To obtain high-resolution, haplotype-specific assemblies of the Sunandini calf (MT431), the TrioCanu module of the Canu assembler was employed. Short-read sequencing of the dam (MT432) and sire (MT433) was performed on the Illumina platform, generating coverages of 31.49× and 33.85×, respectively. After quality trimming and filtering, parental reads were used for trio binning to partition the offspring’s Nanopore long reads into paternal- and maternal-specific sets based on haplotype-specific k-mers (Details of the binned reads are summarized in Supplemental table S5). Using these haplotype-specific read sets, each haplotype was assembled independently with the Long Read Support (beta) plugin of CLC Genomics Workbench v22.0.5. The resulting paternal assembly spanned 2,556,074,938 bp with an N50 of 1.4 Mb, while the maternal assembly measured 2,618,152,939 bp with an N50 of 2.0 Mb (Table 7).

**Table 7.** Summary of Haplotype-Resolved Assembly Statistics.

| Assembly Metric | MT431_Haplotype1 | MT431_Haplotype<br>2 |
| --- | --- | --- |
| Number of contigs | 6,855 | 6,716 |
| Minimum contig length (bp) | 10,218 | 10,230 |
| Maximum contig length (bp) | 1,244,759 | 6,080,185 |
| Average contig length (bp) | 149,064 | 372,343 |
| N50 (bp) | 182,973 | 636,978 |
| N90 (bp) | 83,104 | 171,894 |
| Total assembly length (bp) | 1,021,833,713 | 2,500,658,026 |

### Characterization of genome-wide structural and SNP variation

After comparing to the taurine reference genome ARS-UCD1.2, a total of 16,151 structural variants (SVs) were detected, of which 15,576 were autosomal, ranging in size from 50 bp to 9.94 kbp. These SVs included 5,892 insertions, 4,598 deletions, 1,678 tandem expansions, 1,288 tandem contractions, 1,002 repeat expansions, and 1,693 repeat contractions, collectively affecting 12.79 Mbp of sequence. A Circos plot illustrating each type of SV is shown in Figure 3B. On average, one structural variant was observed every 162,738 bp. Chromosome-wise SV rates, calculated as the number of variants per chromosome length, were highest on chromosome X and lowest on chromosome 26 (Figure 4A). The majority of SVs (∼90%) were located in intergenic and intronic regions, while 232 SVs (∼0.9%) occurred within coding regions (Figure 4B). Among these, 119 were classified as high-impact variants, affecting 78 genes. Predicted functional consequences included frameshift variants, missense variants, start-lost variants, and stop-lost variants. The 78 affected genes were associated with diverse biological pathways, including G protein-coupled receptor activity, response to stress, ubiquitin recycling, and myeloid leukocyte activation. Gene ontology clustering of biological processes is presented in Figure 4C.

**Figure 4.**
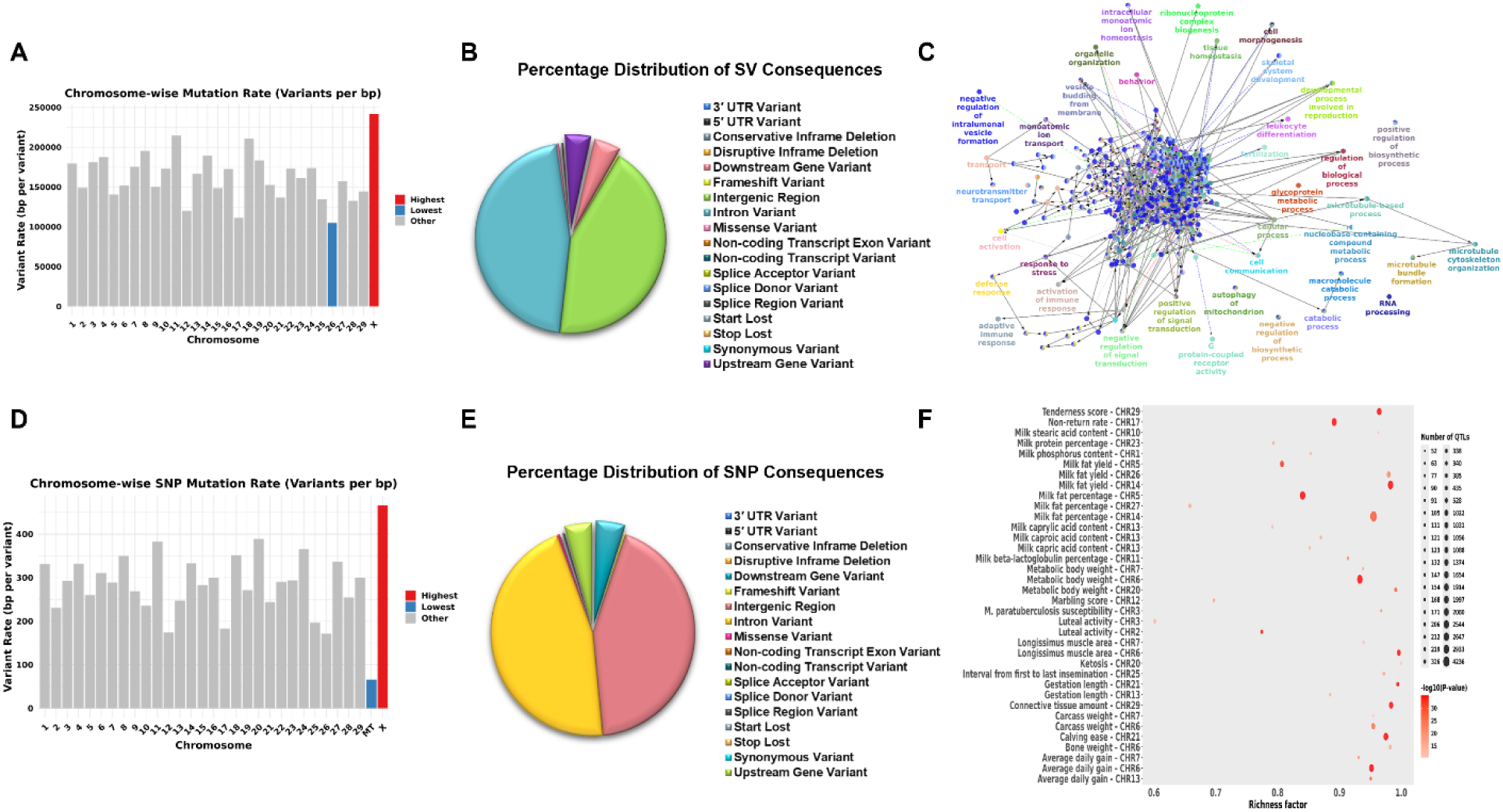
Genome-wide distribution and functional annotation of structural variants and SNPs in the Sunandini genome. (A) Histogram showing the chromosome-wise structural variant rates, with the chromosome exhibiting the highest variant rate highlighted in red and the lowest in blue. (B) Pie chart showing the percentage distribution of structural variants by predicted functional consequence in sample MT431 (calf), as determined using SnpEff. (C) Gene Ontology (GO) analysis illustrating the enriched biological processes associated with genes affected by high-impact structural variants. (D) Histogram showing the chromosome-wise SNP density, with the chromosome exhibiting the highest SNP rate highlighted in red and the lowest in blue. (E) Pie chart showing the percentage distribution of SNPs by predicted functional consequence in sample MT431 (calf), as determined using SnpEff. (F) Bubble plot depicting QTL enrichment analysis for variations that are homozygous in both the dam and sire. Darker red shades indicate higher statistical significance, while the circle area is proportional to the number of associated QTLs. The x-axis represents the richness factor, calculated as the ratio of annotated QTLs to the total number of each QTL type in the reference database.

In addition to structural variants, a total of 9,288,649 single-nucleotide variants (SNVs) were identified. Of these, 8,990,353 were located on autosomes, 298,049 on the X chromosome, and 247 in the mitochondrial genome, corresponding to an average density of one variant every 282 bases, with the highest variant rate observed on chromosome X and the lowest on chromosome 26 (Figure 4D). A total of 143,617 variants (0.99%) were located within coding regions, comprising 50,681 missense variants (0.349%), 71,400 synonymous variants (0.491%), and 96 nonsense mutations (0.001%) (Figure 4E). Functional significance was further assessed through QTL enrichment analysis using the GALLO R package [17], which revealed statistically enriched QTLs associated with key economic traits, including meat tenderness, milk fat yield and percentage, and metabolic body weight (Figure 4F).

### Assessment of genetic relatedness and ancestral origins of Sunandini using genome-wide SNPs

After identifying genomic variants, the ancestral origin of Sunandini was investigated by comparing its variants with a previously published *Bos indicus* Vechur sample and 432 publicly available *Bos taurus* and *Bos indicus* samples from the Bovine Genome Variation Database and the Selective Signatures Database, representing populations from China, Africa, Europe, and India. A total of 19,968,665 high-quality SNPs were used for this analysis. Multidimensional Scaling (MDS) analysis was performed in PLINK to assess genetic relatedness and population stratification. As shown in Figure 5A, Indian *Bos indicus* breeds, including Vechur, clustered together, while Chinese indicine populations formed a separate cluster. Three Sunandini samples clustered closely with European *Bos taurus* breeds, and the remaining two clustered with East Asian *Bos taurus* breeds, indicating a strong taurine genetic background. African populations and Northwest Chinese populations formed distinct clusters but were closer to taurine. These results indicate that Sunandini is genetically closer to *Bos taurus*. Admixture analysis with *Bos indicus* (Brahman, Nellore, Vechur) and *Bos taurus* (Holstein, Hereford, Jersey, Brown Swiss) breeds was performed to further validate this observation (Figure 5B). At both K=2 and higher levels of population subdivision, Sunandini consistently exhibited ancestry components predominantly derived from taurine cattle, confirming a taurine-dominant genomic background.

**Figure 5.**
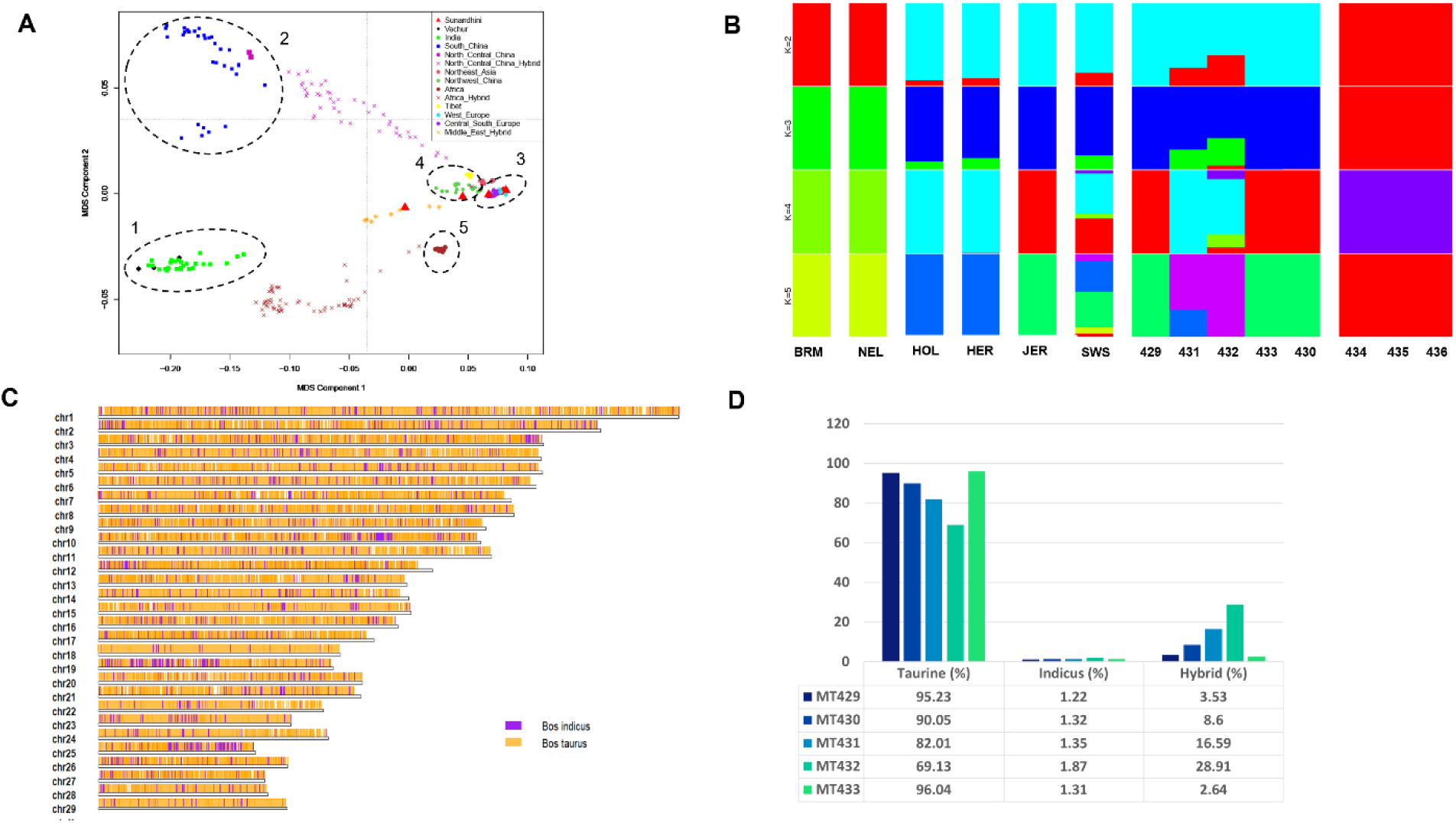
Population structure and genetic relationships among global cattle breeds. (A) Multidimensional scaling (MDS) analysis illustrating the genetic relationships among cattle breeds from different regions worldwide, including Sunandini . The breeds were grouped into five major clusters: Cluster 1 – Indian *Bos indicus*, Cluster 2 – Chinese *Bos indicus*, Cluster 3 – European Bos taurus, Cluster 4 – East Asian Bos taurus, and Cluster 5 – African Bos taurus. Notably, Sunandini clustered with the European Bos taurus group, indicating a closer genetic affinity to taurine cattle. (B) Admixture analysis of 15 cattle breeds, with ancestral clusters ranging from K = 2 to K = 5, illustrating population structure and shared ancestry. The dataset included Brahman (BRM) and Nellore (NEL) representing *Bos indicus*; Vechur (434, 435, 436) representing *Bos indicus*; Holstein (HOL), Hereford (HER), Jersey (JER), and Brown Swiss (SWS) representing Bos taurus; and Sunandini (429, 430, 431, 432, 433) representing crossbred individuals. (C) Karyogram depicting the chromosomal distribution of ancestral genomic segments in Sunandini (sample 432). Orange regions denote *Bos indicus*-derived genomic segments, whereas purple regions represent Bos taurus-derived segments. (D) Bar plot showing the ancestral genomic composition of all five Sunandini samples (429, 430, 431, 432, and 433)

### Tracing taurine and indicine ancestry in Sunandini genome using local ancestry inference

To determine the ancestral composition of genomic regions in Sunandini, local ancestry inference was performed using Lotter software. This method compares each SNP with reference haplotypes from the parental populations (European *Bos taurus* and Indian *Bos indicus*) and probabilistically assigns ancestry along the genome, allowing identification of regions derived from taurine or indicine cattle. For this analysis, 70 million high-quality SNPs from 112 *Bos taurus* and *Bos indicus* samples were used.

The highly admixed sample MT432, identified through admixture analysis, exhibited approximately 69% taurine ancestry, 2% pure indicine ancestry, and nearly 29% hybrid regions with one haplotype taurine and the other indicine. The chromosomal distribution of ancestral genomic segments is depicted in the karyogram in Figure 5C. Analysis of all other individuals revealed a consistent pattern, with the indicine fraction ranging from 1.22% to 1.87%, the taurine fraction ranging from 69.13% to 96.04%, and the hybrid portion ranging from 2.64% to 28.91% (Figure 5D). These results confirm that Sunandini exhibits a predominantly taurine genomic composition with a minor but consistent contribution from indicine ancestry.

The taurine regions, represented by both haplotypes, collectively span 32,599 genes. To evaluate the contribution of taurine ancestry to productivity, milk-yield genes were examined using datasets from CgQTL and CattleQTLdb, which compile results from multiple studies and include 3,076 genes. Remarkably, over 98% (3,023 genes) of milk-yield genes are located within taurine regions (Venn diagram, Figure 6A), suggesting that the strong taurine background substantially contributes to the high milk production observed in Sunandini.

**Figure 6.**
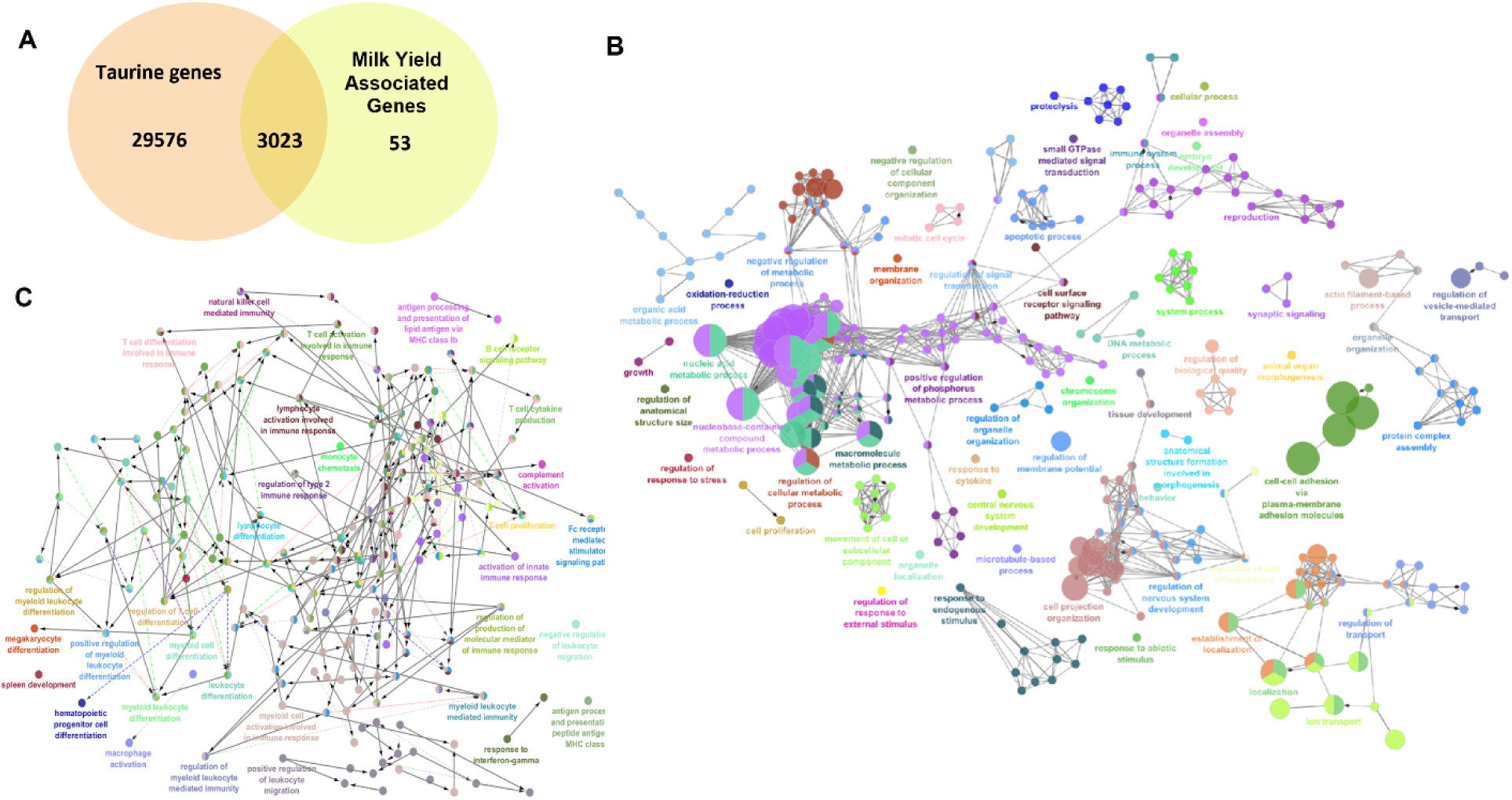
Functional characterization of taurine- and indicine-derived genomic regions in Sunandini cattle. (A) Venn diagram illustrating the overlap between taurine-derived genes identified across all five samples and milk yield–associated genes obtained from the CgQTL and CattleQTLdb databases. (B) Gene Ontology (GO) analysis illustrating the enriched biological processes associated with genes located in the indicine-derived genomic regions across all samples. (C) Gene Ontology (GO) analysis highlighting the enriched immunological pathways associated with genes identified in the indicine-derived genomic regions across all samples.

Within the 1.22-1.87% indicine-derived portion of the genome, approximately 2,931 genes were identified. Gene ontology analysis for biological processes (Figure 6B) revealed that 745 genes are associated with responses to external stress and stimuli, including 11 heat-shock genes. The categories related to stress response and the number of associated genes is listed in Supplemental Table S6. To specifically identify immune-related genes, these were compared with a compiled set of 7,606 genes from InnateDB, ImmPort, the Immunogenetic Related Information Source (IRIS), and the Immunome Database. The intersection of these datasets identified 649 immune-related genes, including 43 key genes involved in innate immunity. These immune genes are enriched in pathways such as T cell differentiation, myeloid leukocyte differentiation, natural killer cell-mediated immunity, antigen presentation, and processing of lipid antigens via MHC class Ib (Figure 6C). Although representing a small fraction of the genome, the indicine-derived regions contribute functionally important traits to Sunandini, notably enhanced stress tolerance and improved immune response.

To evaluate whether highly differentiated regions identified between *Bos indicus* and *Bos taurus* overlap with introgressed regions in Sunandini, heat shock and immune-related genes detected through Fst and XP-CLR analyses were compared with the Sunandini introgression map. Among the heat shock genes, *DNAJB11* (in a taurine region), *DNAJC11*, and *DNAJC5* were located within hybrid regions, indicating that these loci harbor both taurine and indicine haplotypes. For immune-related genes, 32 genes were found in pure indicine regions, whereas 133 genes were located in hybrid regions (summarized in Supplemental table S7). The presence of key heat shock and immune-related genes in hybrid regions suggests that introgressed taurine and indicine haplotypes have jointly contributed to the enhanced thermotolerance and disease resistance observed in Sunandini.

## Discussion

Cattle raised in tropical regions are exposed to constant heat stress and high pathogen pressure. Over time, *Bos indicus* cattle have developed physiological and molecular adaptations that enhance heat tolerance and disease resistance, whereas *Bos taurus* cattle, mainly selected in temperate regions for high milk yield, show lower tolerance to heat and infections. As a result, indicine cattle are better adapted to tropical environments, while taurine cattle perform optimally under favourable conditions. With rising global temperatures, understanding the genetic basis of these differences is essential for developing climate-resilient dairy systems. To address this, the present study compared genome-wide differentiation between *Bos indicus* and *Bos taurus*, and extended the analysis to the tropically adapted crossbred Sunandini population to examine the retention and role of indicine-derived adaptive genomic regions.

Genome-wide sequence-level differentiation was analysed using whole-genome sequencing data from 73 Indian indicine cattle and 84 European taurine cattle. Integration of single-SNP Fst, window-based Fst, and XP-CLR analyses identified 845 highly differentiated loci between the two subspecies. Functional annotation showed that genes involved in cellular stress responses were one of the most strongly differentiated, prompting a focused analysis of heat shock–related genes. Three members of the DnaJ/Hsp40 family, *DNAJC5*, *DNAJC11*, and DNAJC18, were consistently identified by both Fst and XP-CLR analyses, indicating strong and overlapping signals of selection.

*DNAJB11* is an endoplasmic reticulum–localised co-chaperone that interacts with HSPA5 (GRP78) and is involved in protein folding and endoplasmic reticulum–associated degradation (Tan et al. 2014; Behnke et al. 2015). It has been reported to be upregulated under heat stress in bovine granulosa and endometrial cells, as well as in fish, insects, and molluscs (Joyce et al. 2024; Sakai et al. 2022; Wang et al. 2014; Liu et al. 2023; Ge et al. 2024; Ye et al. 2026). *DNAJC5* (cysteine string protein alpha) functions as a co-chaperone of HSPA8 and is involved in preventing protein aggregation, maintaining vesicle stability, and membrane trafficking (Fernández-Chacón and Südhof 2000; Burgoyne and Morgan 2015). It is upregulated in heat-stressed bovine mammary epithelial cells and has been identified within selective sweep regions linked to environmental adaptation in Ladakhi cattle, with similar expression patterns reported in insects and fish (Li et al. 2015; Li et al. 2022; Koloi et al. 2025; Ye et al. 2026). *DNAJC11* is a mitochondrial co-chaperone involved in mitochondrial inner membrane organisation, protein import, and respiratory chain assembly (Xie et al. 2007; Ioakeimidis et al. 2014). It shows differential expression in heat-stressed bovine mammary epithelial cells and has been repeatedly identified in heat-responsive selective sweep analyses of indicine cattle populations (Li et al. 2015; Koloi et al. 2025; Ye et al. 2026; Taye et al. 2018).

Variant annotation of these heat shock genes revealed an enrichment of variants in regulatory regions, splice sites, and untranslated regions, indicating that heat tolerance in *Bos indicus* is largely mediated through regulatory changes rather than protein-coding mutations. Together with their consistent upregulation under heat stress across species, these findings suggest that thermal adaptation occurs mainly through changes in gene expression that help maintain protein folding and mitochondrial function during prolonged heat exposure.

Strong selection signals were also detected in immune-related genes. In total, 235 immune-related genes, including 17 genes involved in innate immunity, showed evidence of positive selection. Gene ontology analysis revealed enrichment of pathways related to lymphocyte activation, T- and B-cell responses, immunoglobulin production, somatic recombination, leukocyte migration, and hematopoiesis, indicating coordinated regulation of adaptive and innate immunity. Variant analysis showed both regulatory and coding differences between *Bos indicus* and *Bos taurus*, suggesting that variation in gene regulation and protein sequence contributes to their contrasting immune responses. Consistent with earlier studies in tick-infested cattle (Rhipicephalus microplus), indicine cattle exhibit stable T-cell mediated adaptive responses, whereas taurine cattle show predominantly innate inflammatory reactions to the same pathogens (Piper et al. 2009; Glass et al. 2005). The overlap of immune loci identified by both Fst and XP-CLR analyses further supports the role of natural selection in immune adaptation to the tropical environment.

To examine how these adaptive features are represented in crossbred cattle, a high-quality, haplotype-resolved genome assembly of Sunandini cattle was generated, showing 95.4% BUSCO completeness. Population structure analyses using multidimensional scaling, admixture, and local ancestry inference revealed a strong predominance of taurine ancestry (69.13-96.04%), with a smaller indicine contribution (1.22-1.87%), reflecting long-term selection for taurine-derived milk production traits.

Consistent with this ancestry pattern, over 98% of milk yield-associated genes (3,023 of 3,076) were located in taurine-derived genomic regions, explaining the high milk productivity of Sunandini cattle. In contrast, although indicine-derived regions constituted only a small fraction of the genome, they were significantly enriched for adaptive genes, including 745 stress-response genes (including 11 heat shock protein genes) and 649 immune-related genes. These genes were mainly associated with pathways such as T-cell receptor signalling, Fc gamma receptor-mediated phagocytosis, natural killer cell-mediated cytotoxicity, leukocyte migration, and phagosome formation, indicating an important role in stress response and immune function under tropical conditions.

Introgression analysis of highly differentiated regions showed that *DNAJC11* and *DNAJC5* were located in hybrid genomic regions, whereas *DNAJB11* was present in a pure taurine region. In addition, 133 immune-related genes were identified in hybrid regions and 32 immune genes in pure indicine regions, suggesting that allele interactions, dosage effects, or novel variants may enhance adaptation. Together, these findings indicate that Sunandini cattle possess a mosaic genome, shaped by the selective retention of advantageous alleles from both taurine and indicine ancestors.

Further validation of these observations using larger populations and functional studies, such as gene expression profiling, will be required to establish direct genotype-phenotype relationships. Such efforts will support the use of adaptive genomic regions in breeding programmes aimed at improving heat tolerance and disease resistance, contributing to sustainable dairy production in tropical and subtropical regions.

## Methods

### DNA isolation from blood samples

Two millilittres of blood samples were collected in a 15-mL Falcon tube, and 4 mL of chilled lysis buffer (150 mM NH4Cl, 10 mM 1M KHCO3, and 0.1 mM EDTA) was added. The samples were gently mixed and incubated on ice for 10 minutes, followed by centrifugation at 7,000 rpm for 10 minutes at 4°C. The supernatant was discarded, and the lysis step was repeated until the pellet was free of red blood cells (two to three washes were sufficient). A total of 300 μL of extraction buffer (400 mM NaCl, 2 mM EDTA, 10 mM TrisCl, pH 8.0) was added and mixed well. Protein digestion was carried out by adding 100 µL of proteinase K (0.2 mg/mL) and 125 µL of 20% SDS, followed by incubation at 56°C for 6 h or overnight. Phenol chloroform extraction was performed by adding 500 µL of phenol–chloroform–isoamyl alcohol (25:24:1) to the mixture, which was gently inverted for 10 min to form a milky emulsion. The mixture was then centrifuged at 10,000 rpm for 6 mins, and the upper aqueous phase was carefully transferred and re-extracted with 500 µL of chloroform: isoamyl alcohol (24:1). DNA was precipitated by adding 1/10th volume of 3M sodium acetate (of an aqueous layer) and 2.5 times volume of chilled absolute ethanol followed by sequential centrifugation at 10,000 rpm for 5 min, 12,000 rpm for 5 min, and 14,000 rpm for 10 min at 4°C. The resulting DNA pellet was washed twice with 300 µL of 70% ice-cold ethanol and air-dried at room temperature. Finally, the DNA was resuspended in 100 µL of nuclease-free water or 1× TE buffer by incubation at 56°C for 10 min and stored at −20°C until further use.

### Genome sequencing and data generation

High-quality genomic DNA was sequenced using both Illumina and Oxford Nanopore platforms. Short-read data generated using the Illumina platform were primarily used for genome size estimation, polishing and error correction of the assembled genome, whereas long reads obtained from the Oxford Nanopore platform were used for the de novo assembly. For Illumina sequencing, the libraries were prepared using the Illumina DNA Prep Kit (Cat. No. 20060060) and sequenced on the NovaSeq 6000 system with an S4 flow cell and NovaSeq 6000 S4 reagent kit v1.5 (300 cycles). In parallel, a long-read sequencing library with an average fragment size of approximately 20 kilobases was prepared according to the manufacturer’s protocol using the Oxford Nanopore Ligation Sequencing Kit and sequenced on the PromethION 24 (P24) platform using FLO-PRO002 R9.4.1 and FLO-PRO112 R10.4 flow cells. Nanopore raw signal data were generated in FAST5 format and subsequently base-called with Guppy to produce demultiplexed FASTQ files containing nucleotide sequences and corresponding quality scores (Q scores).

### De Novo Hybrid genome assembly

The “De Novo Assemble Long Reads” tool within CLC Genomics Workbench (version 22.0.5), supplemented with a specialized plugin for hybrid de novo assembly, was used for genome assembly. This tool is specifically designed to handle long, error-prone reads, such as those generated by Oxford Nanopore Technologies (ONT). It incorporates several open-source assemblers and polishers, including Minimap2, Miniasm, Raven, and Racon. The hybrid assembly workflow comprises two main stages. In the first stage, genome assembly was performed using uncorrected ONT long reads as input, as recommended for this step, and no preprocessing was applied to the nanopore data. The assembly began with the detection of overlap alignment among the input reads using minimap2 and miniasm. The resulting overlaps were processed using pile-o-grams to construct an assembly graph, which was subsequently simplified to generate contigs using the raven assembler. Default parameters were applied, including a k-mer size of 15, a window size of 5, and a minimum contig length of 1,000 bp. Contig quality was further improved using Racon, which employs partial order alignment (POA) of reads against contigs (POA window size = 500 bp) to perform rapid consensus correction. High-quality Illumina short reads were generated by adapter trimming and quality filtering using Trimmomatic v0.39 (Bolger et al. 2014). These processed reads were mapped to the assembled contigs to support further refinement of the genome assembly. The majority of contigs exhibited coverage levels of approximately 40x or higher. Binned reads corresponding to individual contigs were retrieved and used during the polishing process. Two rounds of polishing were performed using racon/minimap2, with a POA window size of 500 bp and a minimum output sequence length of 10,000 bp, as all contigs exceeded this threshold. The polished contigs were scaffolded into pseudo-chromosome-level assemblies using RagTag (v2.1.0) (Alonge et al. 2022) with the *Bos taurus* reference genome ARS-UCD1.2 (GenBank assembly accession: GCF_002263795.1) as a guide under default parameters. A subsequent RagTag patch step was performed using the same reference genome to further improve scaffold continuity. The resulting draft assembly was then subjected to three rounds of polishing using NextPolish (v1.3.127) (Hu et al. 2020). Both quality-filtered Illumina short reads and Oxford Nanopore long reads were used under default parameters to improve overall assembly accuracy.

### Genome assembly quality assessment

The completeness of the final genome assembly was evaluated using BUSCO (version 2.14.1) (Simão et al. 2015) with the mammalia_odb10 dataset, which evaluates gene space completeness based on the presence of conserved single-copy orthologs. To assess mapping consistency and overall assembly quality, Illumina paired-end reads were aligned to the assembled genome using BWA-MEM2. Potential contaminant sequences were identified by examining all unplaced contigs using BLASTn (v2.14.1) (Altschul et al. 1990) by searching against the core_nt database. The BLAST search was performed with the parameters -max_target_seqs 1 -evalue 1e-5 -culling_limit 5, and results were reported in tabular format -outfmt “6 qseqid sseqid pident length mismatch gapopen qstart qend sstart send evalue bitscore qscore sscinames”. Final assembly statistics were generated using Assembly-stats (v1.0.0), and a snail plot was produced with BlobToolKit (Challis et al. 2020) to visualize assembly completeness, quality metrics, and taxonomic composition.

### Haplotype-resolved assembly

Haplotype-resolved assemblies were generated using the TrioCanu module of the Canu assembler (Koren et al. 2018). Prior to assembly, haplotype binning (trio binning) was performed, in which short-read sequencing data from the parental genomes were used to partition long reads from the offspring into haplotype-specific read sets. Each haplotype was then assembled independently, resulting in a complete reconstruction of the diploid genome. For sample MT431, the parental short-read data from MT432 (dam) and MT433 (sire) were quality-trimmed and filtered before being used for trio binning of the long-read data generated from their offspring, MT431. The trio binning divides the total long-read dataset into maternal and paternal read groups based on the presence of haplotype-specific k-mers. These haplotype-specific read sets were subsequently assembled using the Long Read Support (beta) plugin of CLC Genomics Workbench 22.0.5.

### Structural variant analysis

The genome assembly was aligned against the *Bos taurus* (Hereford breed) reference genome ARS-UCD1.2 (GenBank accession: GCF_002263795.1) using NUCmer (Kurtz et al. 2004) with parameters l = 100 and c = 500 to generate a delta file. The resulting alignment file was uploaded to Assemblytics (Nattestad and Schatz 2016) for the identification and analysis of structural variants. The input file (OUT.delta.gz) was provided for loading on the Assemblytics web server, enabling dynamic visualisation and exploration of the results. The functional consequences of the identified structural variants were predicted using SnpEff (Cingolani et al. 2012). Subsequently, pathway analysis of genes with high-impact variants was performed in Cytoscape (Shannon et al. 2003) using Gene Ontology annotations.

### Alignments and single-nucleotide variant identification

Prior to read mapping, adapter sequences and low-quality Illumina reads were removed using Trimmomatic v0.39 (Bolger et al. 2014). High-quality reads were then aligned to the ARS-UCD1.2 bovine reference genome assembly using the BWA-MEM algorithm of the Burrows–Wheeler Aligner (BWA) v0.7.5a with default parameters (Li 2013). Following alignment, SAMtools v1.9 (Danecek et al. 2021) was used to convert SAM files to binary alignment map (BAM) format and to sort mapped reads by chromosomal position. Duplicate reads were identified and removed from the sorted BAM files using the Picard tool’s MarkDuplicates program (v2.17.11). Single-nucleotide polymorphisms (SNPs) were identified using the HaplotypeCaller module of the Genome Analysis Toolkit (GATK) v3.8 (McKenna et al. 2010). All SNPs were subsequently filtered using GATK VariantFiltration with the following threshold criteria: QUAL < 30.0, Quality by Depth (QD) < 2.0, Fisher’s Strand Bias (FS) > 60.0, RMS Mapping Quality (MQ) < 40.0, Strand Odds Ratio (SOR) > 3.0, Mapping Quality Rank Sum Test (MQRankSum) < −12.5, and Read Position Rank Sum Test (ReadPosRankSum) < −8.0.

### QTL enrichment analysis

To investigate the functional and phenotypic implications of genetic variants in Sunandini cattle, quantitative trait locus (QTL) enrichment analysis was performed using the GALLO package in R (Fonseca et al. 2020) on homozygous alternate-allele SNPs identified in both the dam and sire. QTL annotations were retrieved using the find_genes_qtls_around_markers function with a GFF file obtained from The Animal QTL Database (Hu et al. 2019), aligned to the ARS-UCD1.2 reference genome. QTL boundaries were defined as 100 kb upstream and downstream of each significant SNP. Enrichment analysis was conducted by calculating adjusted p-values (Padj) using a chromosome-based false discovery rate (FDR) approach. Traits associated with specific chromosomes with Padj values below 0.05 were considered significantly enriched. Chromosome-level trait enrichment with significant Padj values was visualised using the QTLenrich_plot function.

### Population structure analysis

To validate genetic relatedness and population stratification, the study samples were compared with previously reported SNP data comprising 432 individuals of diverse B. taurus and B. indicus breeds from global populations, which were used as the reference dataset (http://animal.omics.pro/code/index.php/BosVar). The datasets were merged using the vcf-merge tool of VCFtools (Danecek et al. 2011), retaining only SNPs common to both datasets. Classical multidimensional scaling (MDS) analysis was performed in PLINK (v1.07) (Purcell et al. 2007) based on pairwise identity-by-state (IBS) distances, and MDS plots were generated using the R package MDSplot.

For admixture analysis, linkage disequilibrium pruning was performed using PLINK (Purcell et al. 2007) with the parameter --indep-pairwise 50 10 0.1, corresponding to a window size of 50 kb, a step size of 10 bp, and an r² threshold of 0.1 to select a set of approximately independent variants and reduce marker redundancy. The pruned dataset was then analyzed using ADMIXTURE v1.3.0 (Alexander et al. 2009) with default settings and cross-validation (cv = 5), testing the number of ancestral populations (K) from 2 to 5. The admixture results were visualized using custom R scripts.

### Identification and localization of genomic introgressions

Local ancestry inference was performed independently for each chromosome using the Loter v1.0 (Dias-Alves et al. 2018). The analysis used VCF files from two reference populations: 83 *Bos taurus* individuals and 29 *Bos indicus* individuals, along with a third VCF file representing the Sunandini crossbred population. All analyses were conducted using default parameters, with phase correction enabled to improve inference accuracy. The Loter output was used to quantify the genomic proportions and segment lengths inherited from each ancestral population within the Sunandini hybrid genome. The relative contributions of taurine and indicine ancestries across the hybrid genome were visualized using the R package karyoploteR (Gel and Serra, 2017). Genes located within the taurine- and indicine-derived genome segments were annotated, and their biological functions were analysed in Cytoscape (Shannon et al. 2003) using the Gene Ontology dataset to identify enriched functions and associated biological pathways.

### Screening of differentially selected regions

To assess population differentiation between *Bos tauru*s and *Bos indicus*, the fixation index (Fst) was calculated using genome-wide SNP data. Pairwise Fst estimates for autosomal chromosomes were computed using VCFtools (v0.1.13) based on Weir and Cockerham’s method (Danecek et al. 2011; Weir and Cockerham 1984) with default parameters. Both single-locus SNP-based and window-based Fst analyses were performed using a sliding window approach with a window size of 50 kb and a step size of 25 kb to identify broader genomic regions exhibiting population differentiation. In parallel, the Cross Population Composite Likelihood Ratio (XP-CLR) (Chen et al. 2010) method was applied to detect signatures of selective sweeps by integrating multi-locus allele frequency differences and linkage disequilibrium patterns between populations, using the same window and step sizes. Genes located within regions identified by both Fst and XP-CLR were further analysed for functional annotations in Cytoscape (Shannon et al. 2003) using the Gene Ontology dataset to identify enriched biological pathways. Population differentiation signals were visualised using Manhattan plots generated with the qqman R package. (Turner 2018)

### Repeat masking

Repetitive elements in the genome assembly were identified and masked using RepeatMasker v4.1.7-p1 (Smit et al. 2013–2015), a similarity-based tool for detecting interspersed repeats and low-complexity sequences. The genome assemblies were screened against a custom repeat library obtained from Dfam (Dfam-RepeatMasker.lib) (Hubley et al. 2016) using the -lib option. RepeatMasker was run in slow mode (-s) to improve detection of divergent and low-copy repeats, and the -x option was used to generate repeat annotation files in cross_match format. The resulting repeat annotations were subsequently used to produce repeat-masked genome assemblies for downstream gene prediction and functional annotation.

### Genome annotation

Genome annotation was performed for 21 chromosome-level *Bos indicus* and *Bos taurus* genome assemblies using BRAKER3 (Gabriel et al. 2024), an automated gene prediction pipeline that integrates GeneMark-EP/ETP and AUGUSTUS (Stanke et al. 2006) for eukaryotic genomes. Gene prediction was conducted using the default BRAKER3 parameters, with all assemblies provided in soft-masked format and processed with the --softmasking option, which downweights repetitive regions without removing them during gene prediction. BRAKER3 was run in protein-supported mode by supplying a curated vertebrate protein dataset (Vertebrata.fa - https://bioinf.uni-greifswald.de/bioinf/partitioned_odb11/) through the --prot_seq option. These protein sequences were aligned to the target assemblies to generate exon and intron hints and to train AUGUSTUS, enabling accurate gene prediction in the absence of RNA-seq data. Species-specific training was performed automatically within the BRAKER3 workflow to improve gene models for each assembly. Concise

### Macrosynteny analysis

Macrosynteny analysis was performed using the Oxford Dot Plot (ODP) software (Schultz et al. 2023) to assess chromosome-scale conservation of gene order among bovine genomes. ODP is a protein-based synteny framework that compares large and complex genomes across multiple species by using conserved protein-coding genes as anchors. Protein-to-chromosome coordinate files (.chrom) were generated for all genome assemblies as required by ODP. For assemblies annotated at NCBI, .chrom files were generated from GFF3 annotations using the NCBIgff2chrom.py script provided with ODP. For assemblies lacking NCBI annotations, gene models predicted using the BRAKER pipeline were used to generate .chrom files from GTF outputs using a custom R script that followed ODP file format specifications.

Orthologous genes were identified within the ODP pipeline using DIAMOND protein sequence alignments. DIAMOND searches were performed across all species included in the analysis, and orthologs were retained only if they were identified as reciprocal best hits among all species considered. The analysis followed the authors’ recommended ODP settings, including removal of duplicated proteins, to retain only high-confidence orthologs. Conserved syntenic relationships were further examined using ancestral linkage information from the Chordata database provided with ODP.

A custom bovine comparative database was constructed in which the Hereford assembly represented *Bos taurus*, the Nellore assembly represented *Bos indicus*, and Banteng (*Bos javanicus*) was included as an outgroup species. Multi-species reciprocal best-hit and linkage analyses were performed using the odp_nway_rbh workflow, followed by conversion of reciprocal best-hit tables into alignment-format files using odp_rbh_to_alignments. Conserved chromosomal linkages among the bovine genomes were visualized as ribbon plots using the odp_rbh_to_ribbon script.

### Ethics approval and consent to participate

The samples collected for this study involved prior informed consent of the owners of the cattle, and blood samples were collected by veterinarians using the minimum-harm protocols that apply to blood sampling for routine cattle breeding exercise, disease surveillance and diagnostics.

## Data availability statement

All raw and processed sequencing data of the 48 Indian *Bos indicus* generated in this study have been submitted to the Indian nucleotide data archive (INDA) and can be accessed with INDA Accessions number INRP000128 while the sequence data of the five sunandini genomes can be accessed from Indian nucleotide data archive with study ID

## Acknowledgements

We thank Dr. Jose James for his help and encouragement in initiating the Vechur and Sunandini genome sequencing project. Dr. N. Sadananda Singh has been supported by Ramalingaswami fellowship(Department of Biotechnology, India), BT/PR46677 (Department of Biotechnology, India), (DST, India) and IISER TVM research support. Manpreet Kaur was supported by (DST, India). Dr. Rajesh Vakayil Mani, Dr. Balu Bhaskar, Dr. Rajeev Raghavan Pillai and Dr. T. Sajeev Kumar are supported by KLDB. Prof. TV Anilkumar is supported by SCTIMST and IISER TVM. Shivani Kumari and Manpreet Kaur are supported by IISER TVM. Poorvishaa VM is supported by DBT-category I fellowship. Sequencing and initial data analysis were carried out by miBiome, India.

## Author Contributions

Study planning: RVM, BB, RRP, TSK, TVA

Funding: RRP, TSK,SA and NSS

Sample collection:RVM, SA, NSS

Sample processing for Sequencing: RVM, MK, SA, NSS

Data Analysis:PVM, PG, NU, SK, SA, NSS

Manuscript preparation: PVM, NU and NSS

Project co-ordination and Conceptualization: NSS

